# Foam cell induction activates AMPK but uncouples its regulation of autophagy and lysosomal homeostasis

**DOI:** 10.1101/2020.08.13.244640

**Authors:** Nicholas D. LeBlond, Julia R. C. Nunes, Tyler K.T. Smith, Sabrina Robichaud, Suresh Gadde, Marceline Cote, Bruce Kemp, Mireille Ouimet, Morgan D. Fullerton

**Author notes:** To whom correspondences should be addressed: Dr. Morgan Fullerton, Department of Biochemistry, Microbiology and Immunology, Faculty of Medicine, University of Ottawa, 4109A Roger Guindon Hall, 451 Smyth Rd, Ottawa, Ontario, Canada, K1H 8M5.

## Abstract

The dysregulation of macrophage lipid metabolism drives atherosclerosis. AMP-activated protein kinase (AMPK) is a master regulator of cellular energetics and plays essential roles regulating macrophage lipid dynamics. Here, we investigated the consequences of atherogenic lipoprotein-induced foam cell formation on downstream immunometabolic signaling in primary mouse macrophages. A variety of atherogenic low-density lipoproteins (acetylated, oxidized and aggregated forms) activated AMPK signaling in a manner that was in part, due to CD36 and calcium-related signaling. In quiescent macrophages, basal AMPK signaling was crucial for maintaining markers of lysosomal homeostasis, as well as levels of key components in the lysosomal expression and regulation network. Moreover, AMPK activation resulted in targeted up-regulation of members of this network via transcription factor EB. However, in lipid-induced macrophage foam cells, neither basal AMPK signaling nor its activation affected lysosomal-associated programs. These results suggest that while the sum of AMPK signaling in cultured macrophages may be anti-atherogenic, atherosclerotic input dampens the regulatory capacity of AMPK signaling.

## INTRODUCTION

Atherosclerosis precedes and predicts the development of cardiovascular disease, which is a leading cause of morbidity and mortality worldwide. There are numerous risk factors; however, elevated levels of circulating cholesterol (hypercholesterolemia) is a main driver of atherosclerosis. In the early stages, cholesterol-rich low-density lipoproteins (LDL) accumulate in the subendothelial space where it is prone to modifications such as aggregation and oxidation^1–4^. Endothelial activation can subsequently invoke an inflammatory response that stimulates the up-regulation of adhesion molecules and chemoattractant proteins that facilitates the recruitment and transmigration circulating monocytes^5,6^. Within the atherosclerotic microenvironment, monocyte-derived and/or proliferating macrophages play an important role by scavenging modified LDL particles, leading to foam cell formation and propagating a pro-inflammatory plaque milieu^7–10^. Modeling lipid dynamics in cultured macrophages, usually via loading with acetylated-LDL (acLDL), oxidized-LDL (oxLDL) or aggregated-LDL (agLDL), has been crucial to understanding the initiation and progression of atherosclerosis.

AMP-activated protein kinase (AMPK) governs numerous pathways in lipid metabolism and inflammation. Acting as a critical gauge of cellular energy, AMPK inhibits lipid and protein synthesis, while stimulating autophagy and oxidative metabolism^11,12^. AMPK is activated in response to changes in cellular energy and glucose deprivation (via liver kinase B1; LKB1), and to increases in cytosolic calcium (via calmodulin-dependent protein kinase kinase-2; CaMKK2)^13–17^. We and others have linked AMPK to the regulation of macrophage mitochondrial content, fatty acid oxidation, and inflammatory signaling^18–21^. While the contribution of macrophage AMPK to atherosclerosis remains unresolved, AMPK signaling is critical for the differentiation of monocytes to macrophages, as well as the removal of cholesterol from lipid-laden macrophages in culture and *in vivo*^22–24^.

Macroautophagy (herein referred to as autophagy) is an evolutionarily conserved ‘self eating’ process that plays an important role in the processing of lipid-droplet-associated cholesterol prior to its efflux and is important in atherosclerosis progression and regression^25,26^. AMPK is a positive regulator of autophagy and is known to both directly (via phosphorylation of Unc-like 51 kinase; ULK1 and many others) and indirectly (via the inhibition of mechanistic target of rapamycin complex 1; mTORC1) activate autophagy^27,28^. In addition, AMPK has been recently shown to directly promote transcription factor EB (TFEB) transcriptional activity, a master regulator of genes within autophagy and lysosomal biogenesis^29–33^. Interestingly, atherogenic lipids have been shown to both disrupt lysosomal function, while simultaneously stimulating lysosomal biosynthesis and autophagy via TFEB^34^.

As AMPK sits at a nexus of macrophage immunometabolism, we sought to investigate AMPK signaling during in vitro foam cell formation. Here, we report that in response to various forms of modified LDL, AMPK was activated in a CD36- and CaMKK2-dependent manner. Macrophage AMPK signaling was necessary to maintain normal lysosomal function, but unable to stimulate lysosomal biogenesis. However, baseline AMPK signaling was critical for maintaining normal levels of autophagy and lysosomal-associated genes and more importantly, direct AMPK activation significantly augmented total levels of central autophagy components, such as ULK1 and Beclin-1, which was associated with increased nuclear TFEB. Finally, this regulatory input became overwhelmed and failed during foam cell induction, suggesting the importance of the interplay and timeline of AMPK signaling during this process.

## RESULTS

### Atherogenic lipids activate AMPK in bone marrow-derived macrophages

We set out to determine the effect of atherogenic lipids on AMPK signaling in mouse primary bone marrow-derived macrophages (BMDM). Given the tendency to use acLDL to perform *in vitro* macrophage cholesterol experiments, we first assessed AMPK signaling with and without acLDL and observed a significant increase in the activating phosphorylation of AMPK and downstream signaling to a primary AMPK substrate, acetyl-CoA carboxylase (ACC) (Figure 1A).We next used oxLDL and agLDL forms of LDL (Figure 1B, 1C). After 8 and 24 h treatments, AMPK signaling remained active, although not to the same extent as pharmacological activation with the direct allosteric activator A-769662 (Supplementary Figure S1).

**Figure 1.**
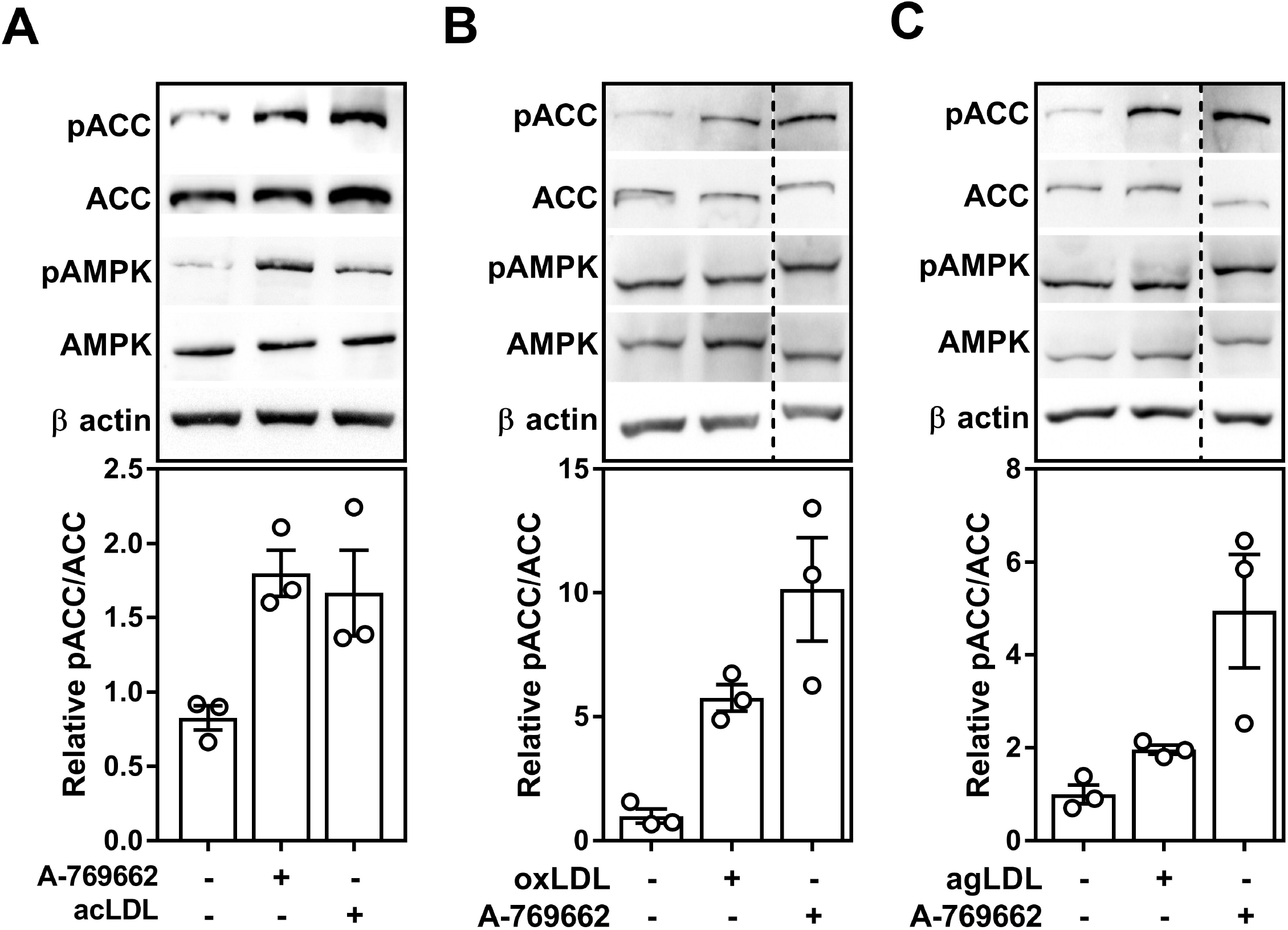
Atherogenic lipoproteins augment AMPK-specific signaling in cultured macrophages. (A-C) BMDM were isolated from WT mice, then incubated with either acLDL (A; 50 μg/mL), oxLDL (B; 50 μg/mL), or agLDL (C; 50 μg/mL) for 24 h (A) and 18 h (B and C). Cells were incubated with A-769662 (A-C; 100 μM) for 24 h (A) and 18 h (B and C) to serve as a positive control for AMPK activation. (A-C) Duplicate gels were used to assess total and phosphorylated proteins. Dashed line signifies where an image was cropped but represent the same gel. Results are representative of 3 independent experiments and data represent the mean ± SEM.

### Atherogenic lipoproteins activate macrophage AMPK partially via CaMKK2

In the context of a plaque-resident, differentiated macrophage, the cause and effect of AMPK activation in response to atherogenic lipids was our next focus. The ER is important for maintaining cytosolic ionic calcium homeostasis and induction of ER stress can lead to increased cytosolic calcium^35^. Atherogenic lipids induce ER stress in BMDM as seen by the increase in IREα and CHOP expression (Figure 2A, 2B), as has been documented in various cell types involved in atherogenesis^3,36^. We hypothesized that uptake and processing of excess cholesterol from atherogenic lipids leads to ER stress and causes activation of AMPK by its upstream kinase, CaMKK2.

**Figure 2.**
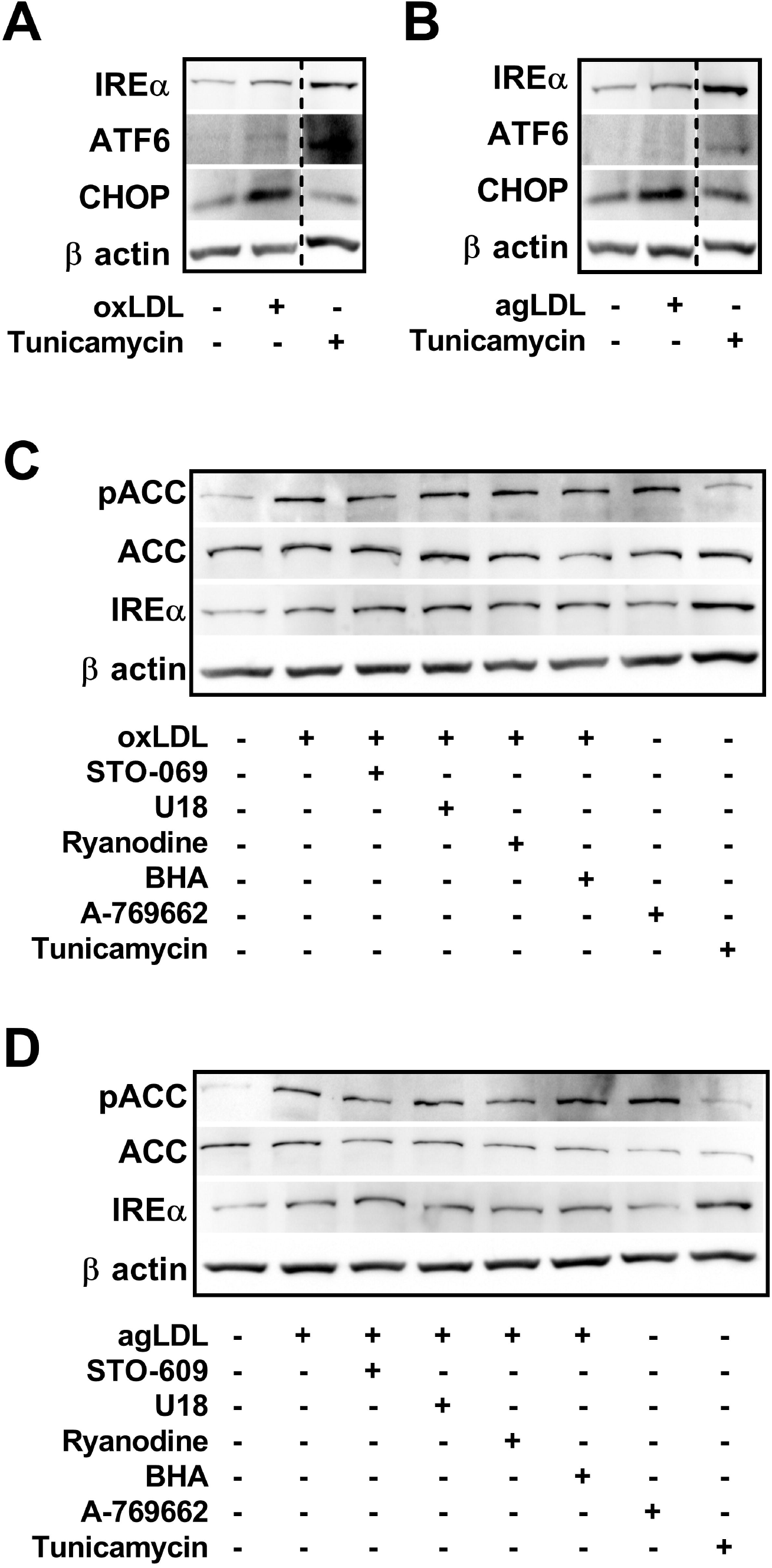
Atherogenic lipoproteins induce modest ER stress and activate AMPK partially through CaMKK2 signaling in cultured macrophages. (A and B) Representative immunoblots depicting the induction of ER stress by up-regulation of its markers in response to oxLDL (A; 50 μg/mL) and agLDL (B; 50 μg/mL). (C and D) Assessing the potential role of CaMKK2 (STO-609; 25 μM), cholesterol trafficking (U18666A; 1 μM), ER-calcium release (Ryanodine; 1 μM), and ROS (BHA; 100 μM) in activating macrophage AMPK in response to atherogenic lipoproteins oxLDL (C) and agLDL (D) through chemical inhibition. (A-D) Macrophages isolated from WT mice were incubated with either oxLDL or agLDL (50 μg/mL) along with inhibitors for 18 h. A-769662 (100 μM) and Tunicamycin (2.5 μg/mL) were used as a positive control for AMPK activation and ER stress, respectively. Data is representative of 3 independent experiments using BMDM isolated from separate mice.

To address this, we co-cultured BMDM with atherogenic lipids in combination with inhibitors of cholesterol trafficking from the lysosome (U18666A), ER calcium release (Ryanodine), ROS generation (BHA), or CaMKK2 (STO-609), and measured AMPK-specific signaling to ACC. Tunicamycin and A-769662 were used as positive controls for ER stress and AMPK activation, respectively. AMPK signaling to ACC was stimulated in the presence of both oxLDL and agLDL. This was partially attenuated by the direct inhibition of CaMKK2 by STO-609 and by inhibition of calcium release by Ryanodine (Figure 2C, 2D). These effects were more pronounced in the agLDL-treated macrophages. Neither U18666A, nor BHA treatment altered AMPK signaling. Interestingly, while both oxLDL and agLDL treatments induced mild ER stress (increased expression of IREα), ER stress induction via tunicamycin did not result in augmented signaling to ACC; however, AMPK activation via A-769662 showed a trend toward improving ER stress.

LKB1 regulates AMPK activity in response to changes in adenine nucleotide ratios. We measured the levels of AMP and ATP in BMDM incubated with atherogenic lipids and observed a consistent but not statistically significant increase in AMP: ATP ratio, measured by HPLC (Figure 3). Taken together, these results suggest that calcium/CaMKK2 and potentially changes in energy, but not ROS, play a role in mediating lipid-induced AMPK activation.

**Figure 3.**
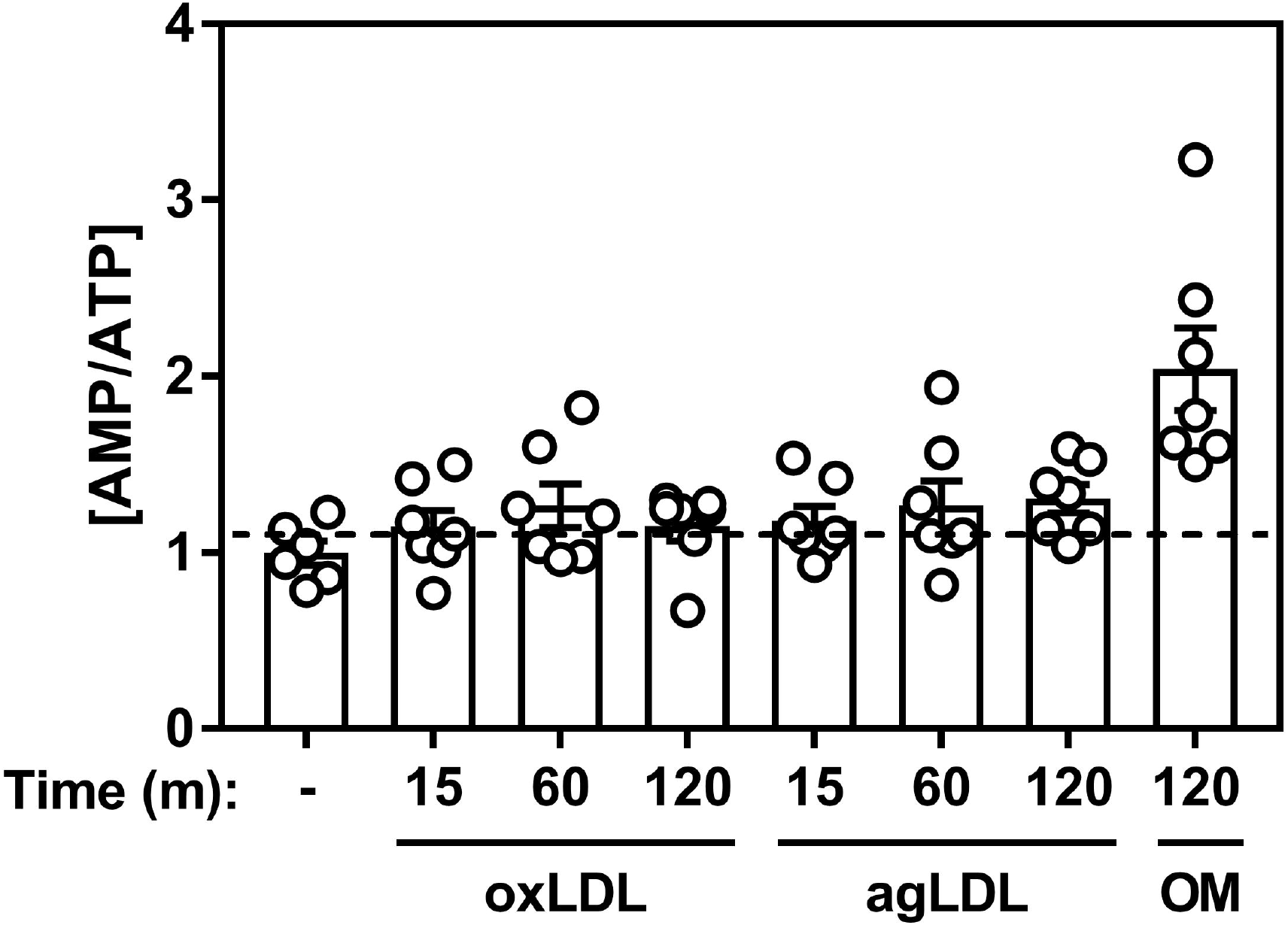
Acute incubation with atherogenic lipoproteins minimally shifts the energy status in cultured macrophages. Macrophages were cultured in the presence of oxLDL (50 μg/mL), agLDL (50 μg/mL), or oligomycin (5 μM) as a positive control. AMP and ATP were measured by high performance liquid chromatography. Data represent the mean ± SEM and expressed relative to naïve BMDM control.

### CD36 links atherogenic lipids, AMPK signaling and autophagy

Autophagy is a highly conserved and regulated process that has been shown to be important for mobilizing cholesterol from macrophage foam cells^37^. There is evidence that upon lipid-loading of cultured macrophages, autophagy is initiated and bulk programs that control autophagy and lysosomal biogenesis are stimulated^34^. To link the observation that atherogenic lipids activated AMPK in macrophages, we treated WT and AMPKβ1-deficient macrophages with acLDL and assessed markers of autophagy. Importantly, as we have shown previously, AMPKβ1-null macrophages had a dramatic (~90%) reduction in AMPK signaling to ACC or ULK1, either in response to AMPK-specific activation (A-769662) or acLDL treatment (Figure 4A). To further investigate AMPK-dependent autophagy signaling, we treated WT and AMPKβ1-deficient macrophages with and without chloroquine (to inhibit autophagosome and lysosomal fusion), A-769662 (to activate AMPK) and acLDL (as an atherogenic stimulus) (Figure 4A). Compared to WT control cells, AMPK deficiency did not alter autophagic flux as seen by the conversion of LC3I to LC3II in the presence of chloroquine. Interestingly, in response to acLDL-mediated lipid loading, LC3II is similarly augmented in both WT and AMPKβ1-null cells (Figure 4B).

**Figure 4.**
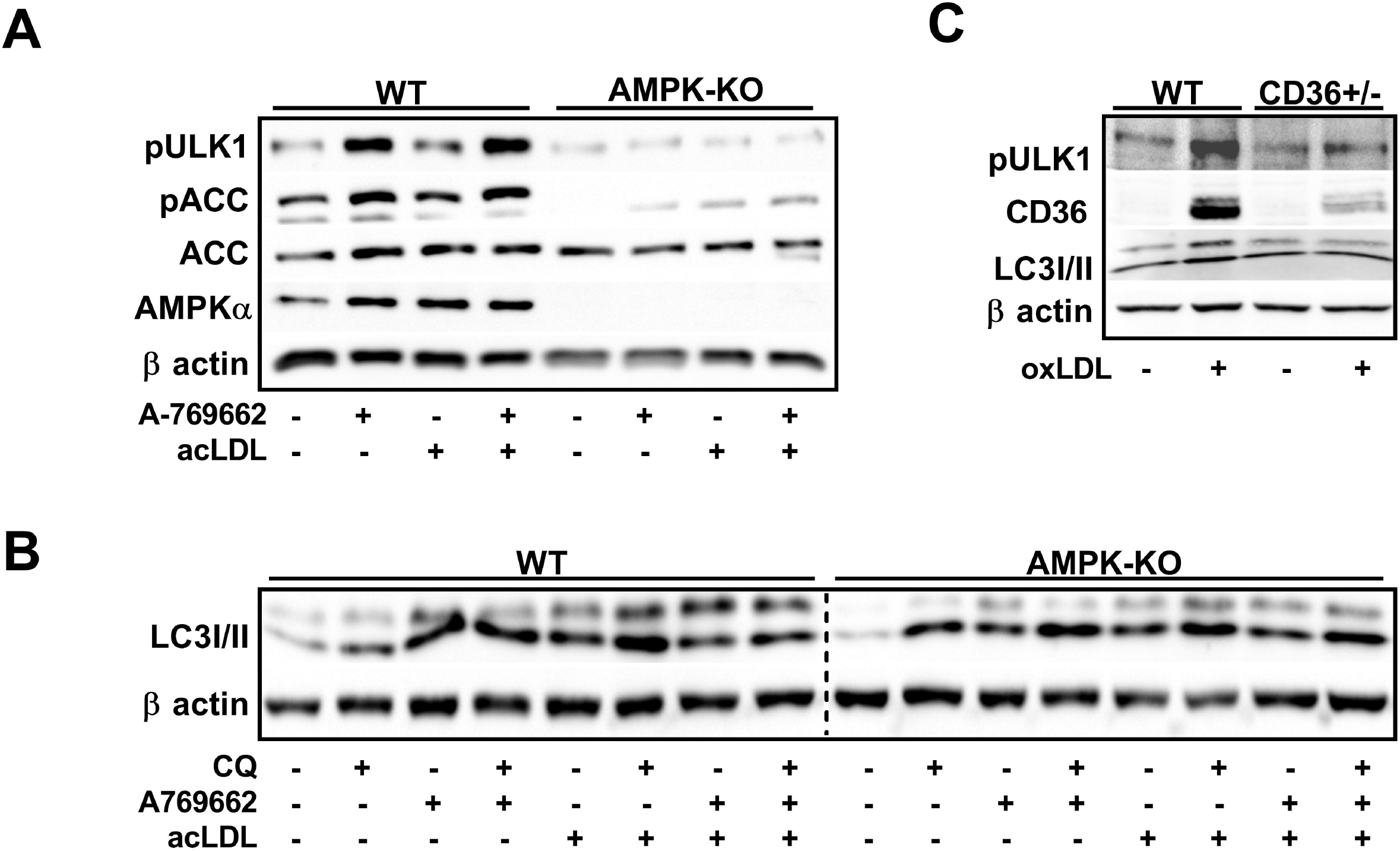
CD36 transmits lipid-induced AMPK activation signal in cultured macrophages. (A) Cells were treated ± acLDL (50 μg/mL) and ± A-769662 (100 μM) for 24 h prior to harvest. (B) Macrophages were treated ± AMPK activation (A-769662; 100 μM), autophagy inhibition (chloroquine; 25 μM), and atherogenic lipoproteins (acLDL; 50 μg/mL). (C) WT and CD36-deficient macrophages were incubated with oxLDL for 24 h (50 μg/mL). (A and C) Total and phosphorylated proteins were determined from duplicate gels. Results are representative of 3 independent experiments from separate mice.

The scavenger receptor CD36 plays a pivotal role in the recognition and unregulated uptake of atherogenic lipids such as oxLDL^7^. There is also evidence that AMPK regulates CD36 and vice versa^38,39^. We next questioned whether CD36-mediated the activation of AMPK signaling in response to atherogenic lipids^39,40^. Treatment with oxLDL dramatically enhanced AMPK-specific ULK1 Ser555 phosphorylation and increased the conversion of LC3II in WT, but not CD36^+/-^ cells (Figure 4C), suggesting that CD36 plays a role in transmitting the atherogenic signal to AMPK, which in turn signals to regulate autophagy programs.

### Macrophage AMPK regulates autophagy signaling

Independent of atherogenic lipid-loading, we were interested in how macrophage AMPK signals to regulate autophagy. As above, the lipidation of LC3I to LC3II and the protein content of p62 were increased in the presence of chloroquine in both WT and AMPKβ1-null cells (Figure 5). However, activation of AMPK did not reveal changes in autophagy flux that were AMPK-dependent. Therefore, we focused on upstream signaling pathways. In WT cells, AMPK activation increased the phosphorylation of ULK1 at Ser555 (an AMPK-specific site), without altering the levels of ULK1 Ser757 (an mTORC1-specific site) phosphorylation. Moreover, activation of AMPK was associated with slight increases in the phosphorylation of RAPTOR and TSC2 in WT, but not AMPKβ1-null cells (Figure 5), which are classically known to facilitate mTORC1 inhibition. However, in AMPKβ1-deficient macrophages, where mTORC1 inhibition of autophagy might be expected to prevail, ULK1 Ser757 phosphorylation was lower when compared to WT control cells. Interestingly, AMPK activation led to a potent up-regulation of ULK1 protein content, whereas there was a dramatic reduction in ULK1 in AMPKβ1-null cells compared to control. Chloroquine treatment also augmented ULK1 protein content; however, this was independent of AMPK signaling.

**Figure 5.**
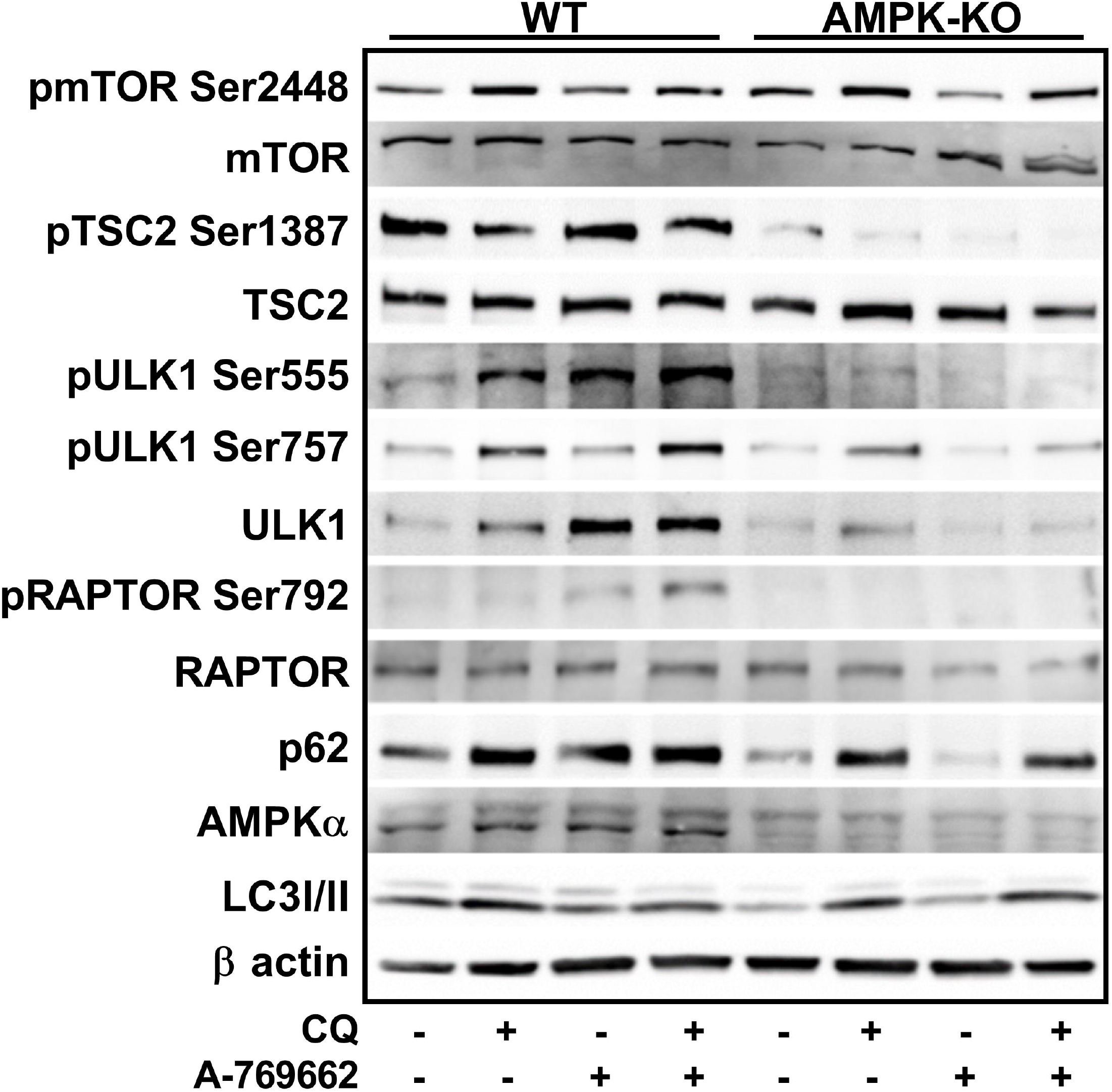
Macrophage AMPK signals through autophagy and is linked to autophagic protein expression. Signaling and flux through autophagy in response to AMPK expression and activation (A-769662 100 μM) and/or autophagy inhibition (chloroquine 50 μM) for 5 h. Total and phosphorylated proteins were determined from duplicate gels. Data is representative of at least 3 independent experiments from BMDM isolated from separate mice.

### Transcriptional control of lysosomal programs by macrophage AMPK is TFEB-dependent

It is known that AMPK signals to ULK1 (at various sites)^27,41^ to initiate autophagy. While we confirmed in macrophages that there was a down-regulation of ULK1 expression in AMPK-deficient cells^29,24^, we also observed a consistent up-regulation of ULK1 protein in response to direct AMPK activation (Figure 5). Given the recent link between TFEB and AMPK, we queried whether acute activation of AMPK altered the transcript expression of several lysosomal and autophagic genes. Transcript expression of TFEB and regulated downstream genes Beclin-1, lysosomal acid lipase (LAL) and LC3 were all lower in AMPKβ1-deficient macrophages compared to WT control cells (Figure 6A-D). Over the course of 24 h, A-769662 WT-treated cells experienced a greater up-regulation of these targets, whereas AMPKβ1-null cells remained mainly unchanged. We then confirmed that the protein expressions of TFEB-regulated ULK1 and Beclin-1 were significantly reduced in AM PKβ ‘-deficient BMDM, compared to WT control, and that, A-769662 increased the total protein expression of both ULK1 and Beclin-1 in macrophages from WT mice but not AMPKβ1-null cells (Figure 6E). Given that TFEB is implicated in the regulation of these targets, we treated macrophages with A-769662 to activate AMPK and observed a dramatic shift in the nuclear enrichment of TFEB (Figure 7A, 7B). However, when TFEB mRNA levels were silenced, the transcript expression of downstream targets ULK1, Beclin-1 and p62 that were associated with A-769662-mediated AMPK activation remained unresponsive (Figure 7D, 7E and Supplementary Figure S2). Taken together, this suggests that in macrophages, AMPK activation signals via TFEB to control lysosomal programs.

**Figure 6.**
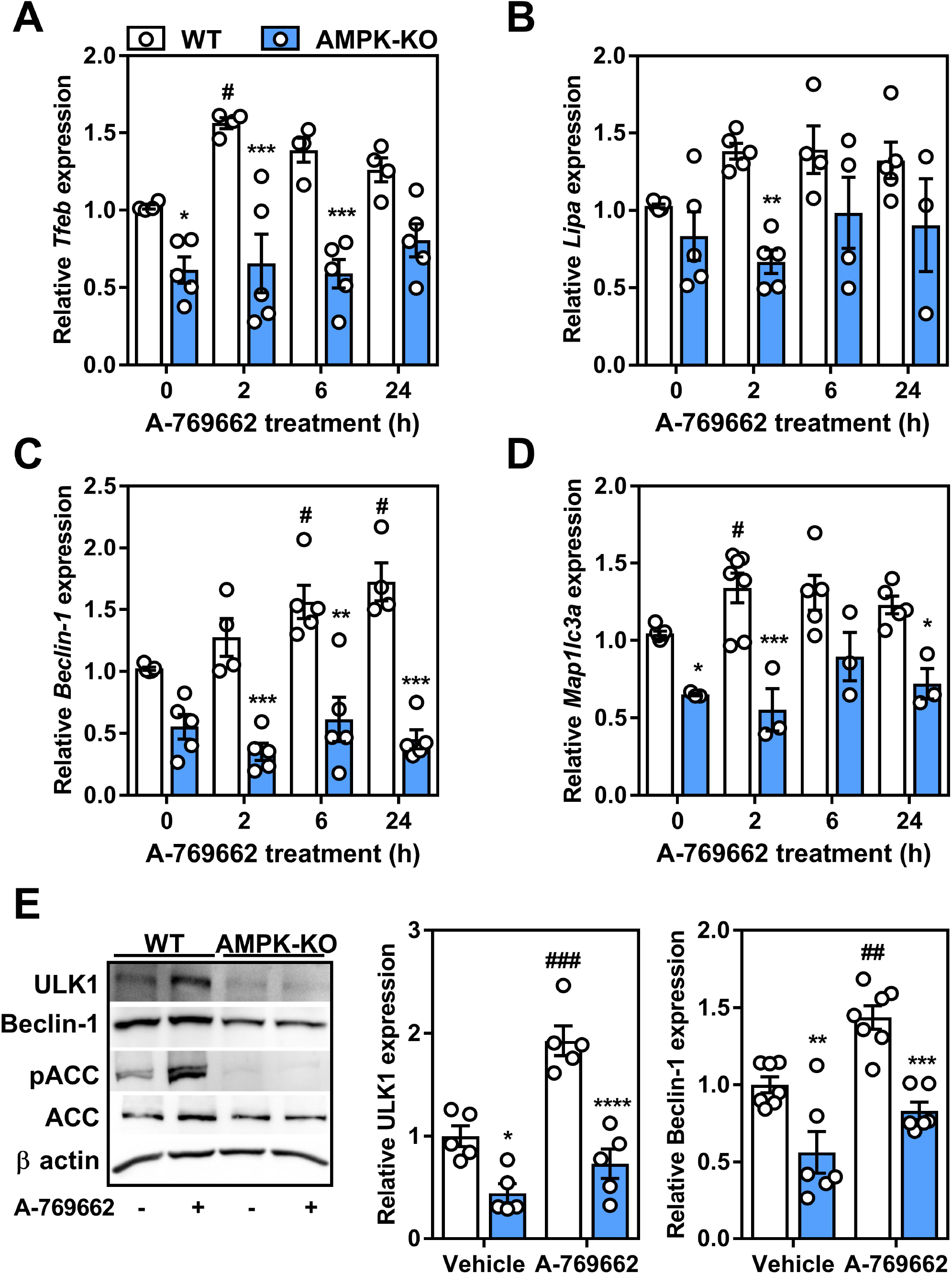
Macrophage AMPK activation leads to the upregulation of autophagy-related genes. Relative transcript expression in response to AMPK activation (A-769662 100 μM) over time for (A) TFEB (*Tfeb*), (B) lysosomal acid lipase (*Lipa*), (C) Beclin-1 (*Beclin-1*), and (D) LC3 (*Map1lc3a*). Transcript expression was normalized to the average expression of *β-actin* and *Tbp*. (E) Protein expression of ULK1 and Beclin-1 were determined following treatment ± A-769662 (100 μM) for 24 h. Data represent the mean ± SEM, where *, **, *** and **** represent p<0.05, p<0.01, p<0.001 and p<0.0001 between genotypes and where ^#^, ^##^ and ^###^ represents p<0.05, p<0.01 and p<0.001 between treatments, as calculated by a 2way ANOVA.

**Figure 7.**
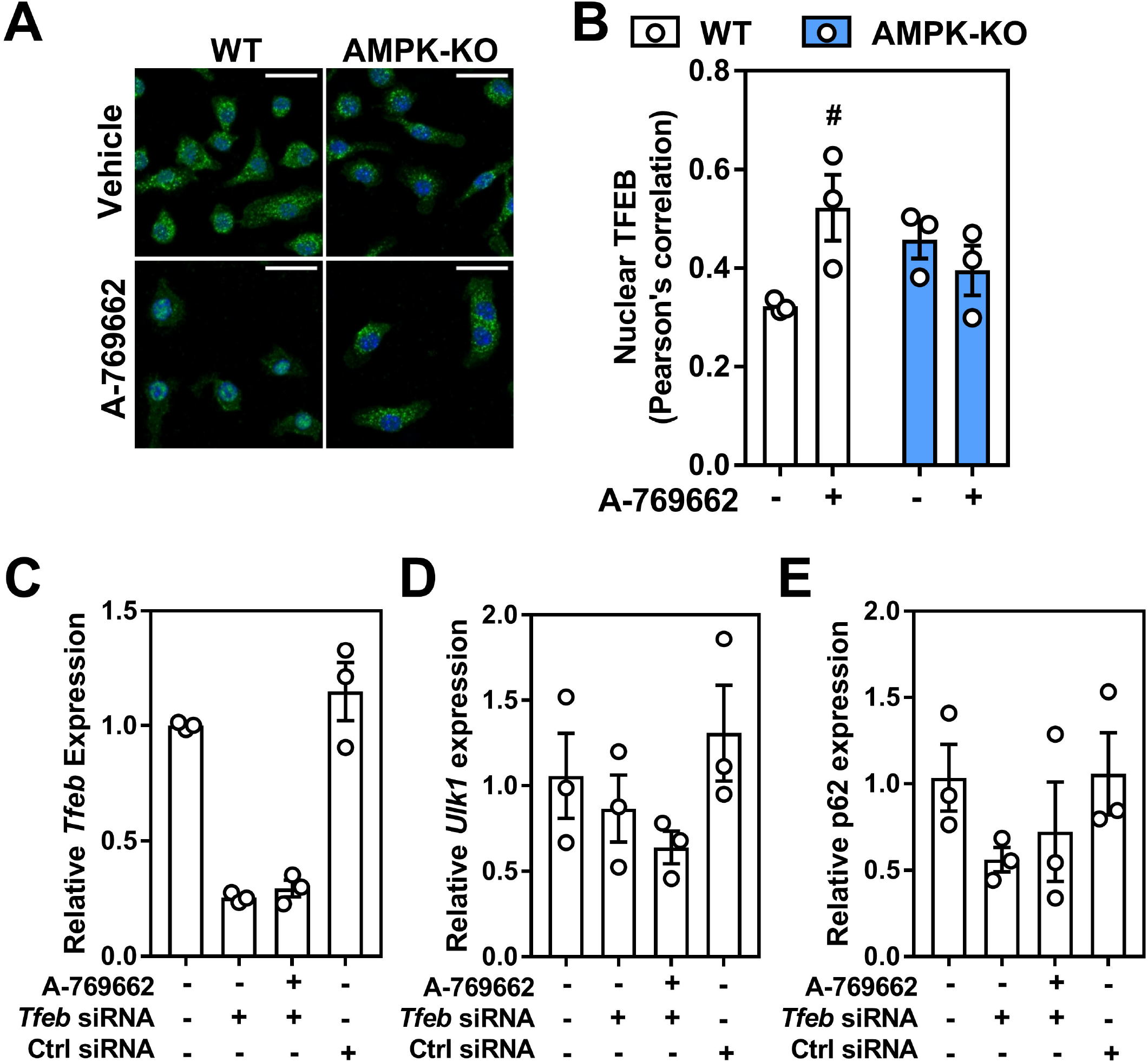
Macrophage AMPK activation regulates lysosomal-associated gene transcription through TFEB regulation. (A) Macrophages were treated for 2 h with A-769662 (100 μM). Shown is a representative immunofluorescent image z-projection for maximum intensity, scale bar is 25 μm (TFEB; Green, DAPI; Blue) (B) Pearson’s correlation coefficient for the co-localization of TFEB and DAPI across all z-stacks from (A), where 1 represents 100% co-localization and 0 represents 100% no co-localization. Macrophages were incubated with *Tfeb* or control siRNA followed by AMPK activation with 5 h incubation with A-769662 (100 μM). The transcript expressions of (C) *Tfeb*, (D) *Ulk1* and (E) *Sqstm1* (gene encoding p62). Transcript expression was normalized to the average expression of *β-actin* and *Tbp*. Data represent the mean ± SEM, where ^#^ represents p<0.05 between treatments, as calculated by a 2way ANOVA.

### Lysosomal programs are partly regulated by AMPK

Since AMPK signaling played a role in controlling transcript levels of important autophagy and lysosomal components, we next determined functional parameters related to lysosomal function in BMDM from WT and AMPKβ1-deficient mice. While lysosomal homeostasis is a highly dynamic process, AMPK deficiency was associated with a significant reduction in the overall number of acidic compartments (taken to broadly represent lysosomal and endosomal compartments), while trending toward a lower ability to cleave a set substrate (interpreted as proteolytic cleavage) compared to WT control cells (Figure 8A and B). There were no differences in measures of lysosomal membrane permeability (Figure 8C). While representing static measures, these results support a broad role for AMPK signaling in facilitating lysosomal programs. Finally, we assessed if these functional measures of lysosomal homeostasis were augmented with AMPK activation. Surprisingly, AMPK activation via A-769662 had no effect on total cellular acidity or total LAMP1 staining and a suppressive effect on proteolytic cleavage (Figure 8D-F).

**Figure 8.**
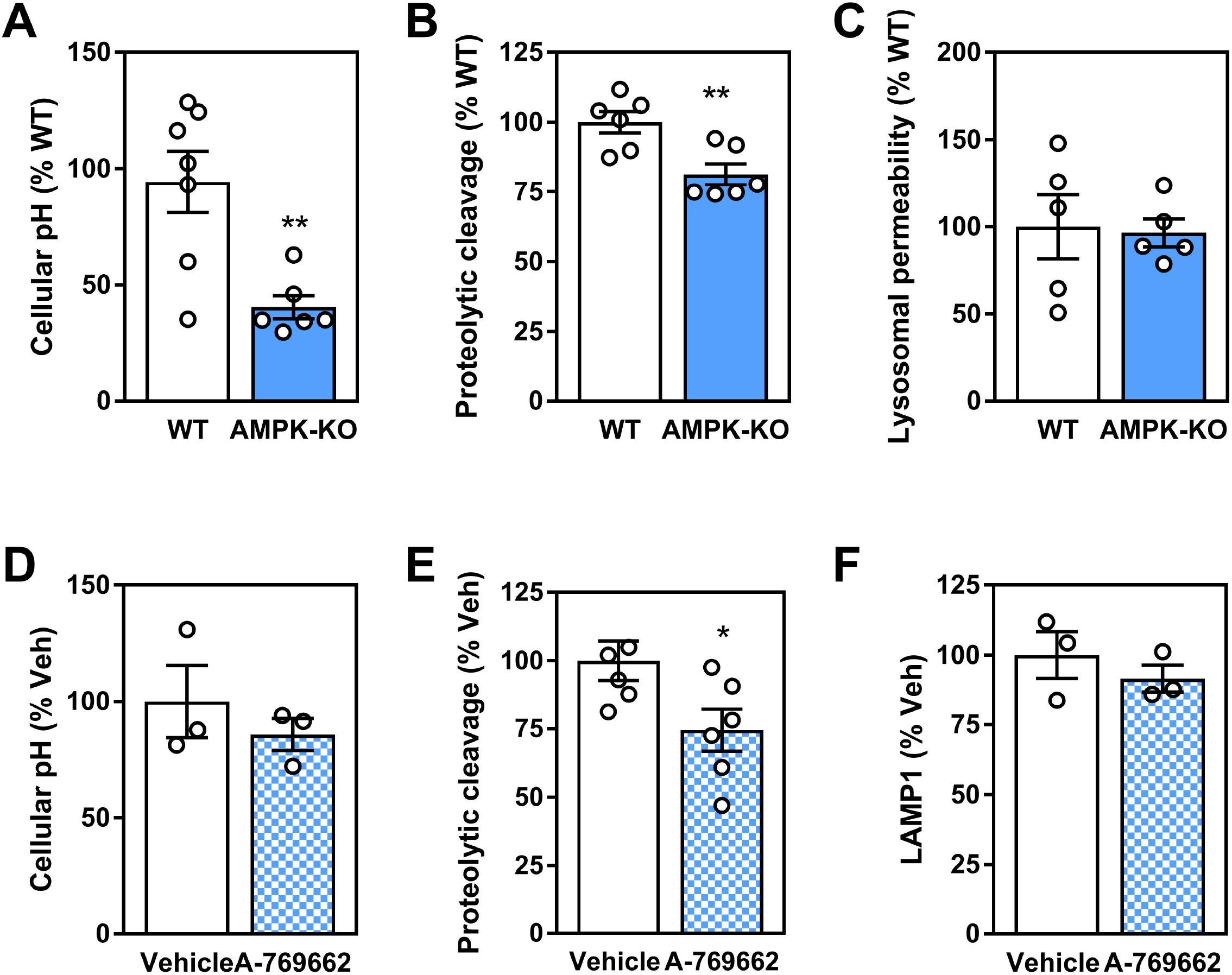
Macrophage AMPK expression and activation influences parameters for lysosomal homeostasis. WT and AMPKβ1-deficient macrophage were assessed for (A) cellular acidity, (B) DQ-ovalbumin proteolytic cleavage and (C) lysosomal membrane permeability. WT and AMPKβ1-deficient macrophages were treated ± A-769662 (100 μM) for 24 h followed by measurement of (D) cellular acidity (E) DQ-ovalbumin proteolytic cleavage and (E) LAMP1 staining. Data represents the mean ± SEM, where * and ** represent p<0.05 and p<0.01 as calculated by two-tailed Student’s t-test.

### Atherogenic lipids overwhelm AMPK regulation of lysosomal programs

It has been shown that atherogenic lipids acutely stimulate a transcriptional lysosomal program^34^. Given our data that atherogenic lipids activate AMPK and that AMPK activation can influence TFEB-regulated lysosomal programs, we hypothesized that AMPK is responsible for the lipid-induced stimulation of lysosomal pathways. Using BMDM, we incubated WT and AMPKβ1-null cells with atherogenic lipoproteins for 3-12 h and assessed the transcript expression of TFEB and two downstream effector genes. Unexpectedly, we observed no induction of TFEB, Beclin-1, or LAL transcript by either oxLDL or agLDL (Supplementary Figure S2). Moreover, in oxLDL- or agLDL-derived foam cells, A-769662 had no effect on TFEB-related transcript levels (Figure 9A-C). Complementary to this, upon oxLDL treatment, no differences were observed in the amount of TFEB in the nucleus. Finally, when cells were primed with atherogenic lipids, AMPK signaling and regulation of lysosomal function was largely uncoupled (Figure 9E-F).

**Figure 9.**
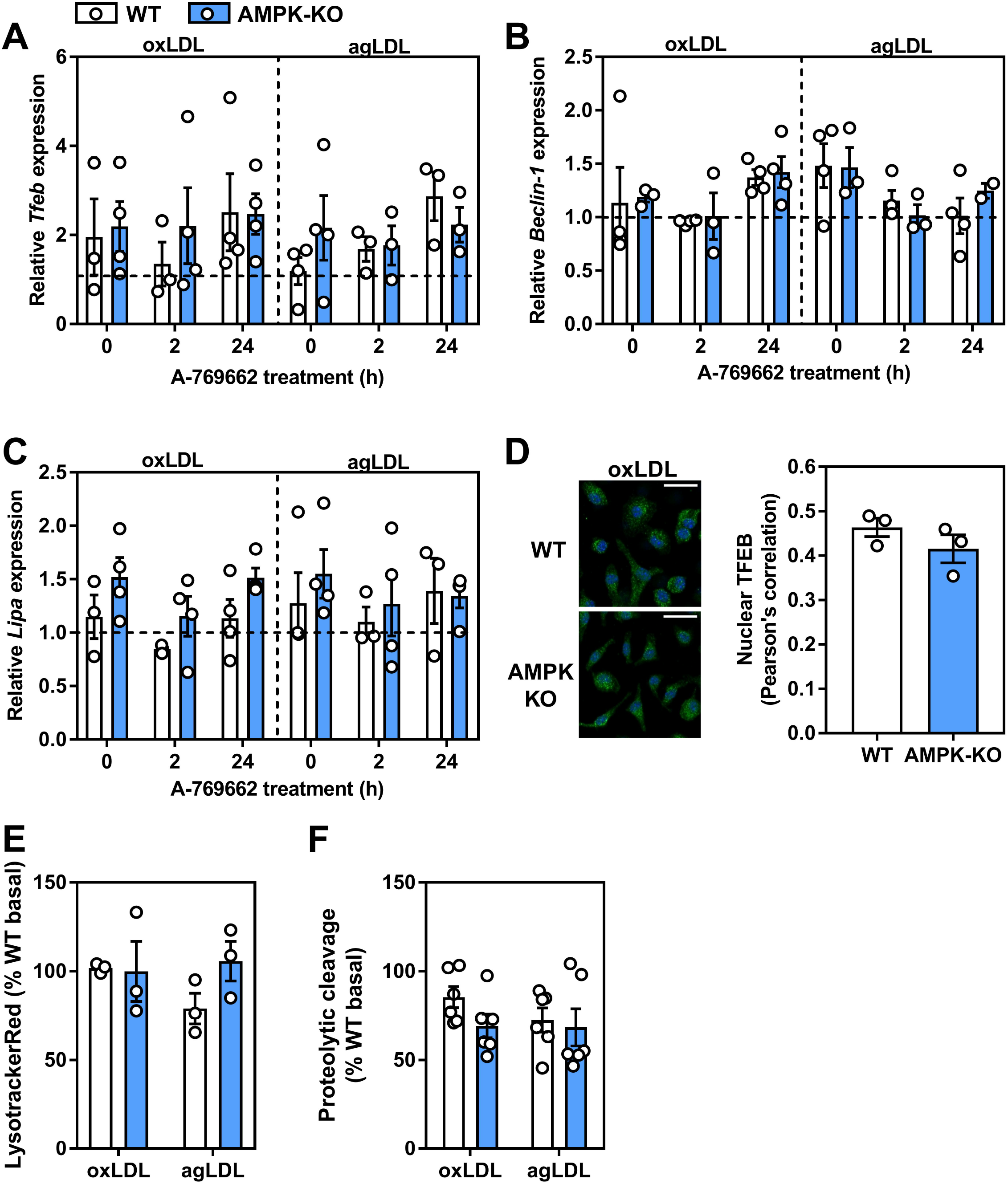
Atherogenic lipoproteins uncouple AMPK from its regulation of lysosomal-associated gene transcription. WT and AMPKβ1-deficient macrophages were cultured for 48 h in ox-LDL- or agLDL-containing media (50 μg/mL each), followed by a 24 h-treatment with A-769662 (100 μM). Relative transcript expression of (A) *Tfeb*, (B) *Beclin-1* and (C) *Lipa* (lysosomal acid lipase) was determined by normalizing to the average of *β actin* and *Tbp*. The horizontal hashed line represents the expression of untreated WT BMDM. (D) WT and AMPKβ1 knockout cells were treated with oxLDL (2 h; 50 μg/mL) and imaged. Representative immunofluorescent image z-projection for maximum intensity, scale bar is 25 μm (TFEB; Green, DAPI; Blue). Pearson’s correlation coefficient quantification of the entire z-stack (right). WT and AMPKβ1-deficient cells were co-cultured with oxLDL and agLDL (50 μg/mL) for 24 h followed by determination of (E) Lysotracker Red staining and (F) DQ-ovalbumin proteolytic cleavage. Data represent the mean ± SEM.

## DISCUSSION

The unregulated uptake and retention of cholesterol-rich lipoproteins is a hallmark of atherogenesis^10,42,43^, yet a complete understanding of the molecular mechanisms that regulate these processes is still lacking. In this study, we aimed to determine if AMPK signaling was affected by treatment with atherogenic lipoproteins and if so, the potential mechanism(s) and consequences. Using bone marrow macrophages, we demonstrated that AMPK signaling was stimulated in response to various types of atherogenic lipoproteins via the calcium activated CaMKK2. However, although AMPK is important in macrophages for the induction and maintenance of lysosomal and autophagic pathways (TFEB), treatment with atherogenic lipoproteins negates these effects.

AMPK signaling is required for proper monocyte-macrophage differentiation and in the human THP-1 monocyte cell line, as exposure to differentiating stimuli (including oxLDL, phorbol esters and 7-ketocholetsterol), resulted in AMPK activation^24^. To our knowledge, our studies are the first to investigate AMPK signaling in response to model (acLDL) and physiological (oxLDL/agLDL) atherogenic stimuli in BMDM. We first focused on a potential mechanism to explain how exposure to these modified lipoproteins could lead to AMPK activation, which can occur in response to glucose deprivation (lysosomal activation), adenine nucleotide fluctuations (AMP/ADP binding to the γ subunit of AMPK), intracellular calcium release (via CaMKK2) and most recently binding of long chain acyl-CoA^44^. Since it has been shown that oxLDL is capable of triggering ER stress, a process linked to alterations in cytosolic calcium flux^3,35,36^, we reasoned that calcium-stimulated CaMKK2 may be responsible. Using tunicamycin and A-769662 as positive controls for ER stress and AMPK activation respectively, oxLDL and agLDL treatments caused a mild ER stress response. Since tunicamycin-induced ER stress did not affect AMPK signaling, it would be misleading to conclude that ER stress itself was responsible; however, in lipid-loaded macrophages, it appears that AMPK signaling is largely affected by the release of calcium and the upstream activation of CaMKK2.

Energy fluctuations are reflected by changes in AMP:ADP:ATP ratios. Upon treatment with atherogenic lipoproteins, there was a small trend toward an increase in AMP: ATP ratios (Figure 3). The activity of the upstream kinase LKB1 had been described as constitutively active, making its phosphorylation and activation of AMPK a function of nucleotide binding and AMPK conformation^15^. Our results do not rule out the potential for LKB1-mediated AMPK activation, which may be responsive to changes in energy metabolism upon lipid loading^45^. Interestingly, binding of long chain acyl-CoA molecules to a conserved region of the AMPK β1 subunit causes allosteric activation (providing endogenous allosteric activation). Therefore, it remains entirely plausible that fatty acids from modified LDL metabolized in a manner that facilitates the allosteric activation of AMPK. Finally, AMPK activation in response to glucose deprivation has been shown to be intricately tied to glycolytic flux (fructose 1,6 bisphosphate binding to aldolase being the key step) and the lysosomal localization of the AMPK-LKB1-Axin activating complex^46^. The physiological relevance of this activation pathway is still being evaluated and provides the basis for exploring whether plaque-resident cells have sufficient access to glucose during atherosclerosis. However, in our study, BMDM were cultured in a surplus of glucose in the media (25 mM), making this an unlikely contributing mechanism for AMPK activation.

Macrophages undergo a classic up-regulation of scavenger receptors upon differentiation from monocytes, which in turn facilitate the unregulated uptake of modified lipoproteins. CD36 is an important receptor that has been implicated in hepatic AMPK signaling^39^. We wondered if in cultured macrophages, CD36 transmitted the lipid-associated signal that caused AMPK activation. We observed that CD36 deficiency dramatically reduced the phosphorylation of ULK1 and the conversion of LC3-I to LC3-II, indicating that CD36 expression controls downstream signals to AMPK and subsequently, autophagy.

Our results indicate that autophagy is active in macrophages even in the absence of AMPK signaling. However, this is where complexities in autophagic flux interpretation come into play. While we conclude that autophagic flux is maintained in both WT and AMPKβ1-deficient macrophages, AMPK activation resulted in more LC3 lipidation in WT cells, such that when autophagy was inhibited with chloroquine, there were no observable differences in basal or lipid-loaded conditions (Figure 4B). Moreover, chloroquine treatment in AMPKβ1-null macrophages showed a consistent increase in p62 protein content that was basally lower compared to WT control cells and not up-regulated in response to AMPK activation (Figure 5). Therefore, our data show that AMPK signaling increases the expression of autophagic machinery and potentially its capacity, but the limitations with measuring bulk autophagic flux by immunoblotting methods prevents further interpretation. Future work is warranted using more autophagy-specific tools^47^.

Our understanding of the role of AMPK-specific signaling in autophagy is continually expanding. While most studies have focused on acute, phospho-regulation of downstream targets, a clearer picture of transcriptional regulation is beginning to emerge. TFEB, a master regulator of lysosomal and autophagic programs, has recently been shown to be regulated both directly and indirectly by AMPK^29,32,33^. Our data suggest that normal AMPK signaling is required in murine macrophages to maintain regular transcriptional and cellular control over lysosomal programs. Since the discovery of AMPK signaling to ULK1, there has been little focus on potential transcriptional control of ULK1 in response to AMPK signaling. Our data corroborates recent work demonstrating that in myeloid cells, levels of ULK1 and Beclin-1 are drastically diminished when AMPK expression is restricted^24,48^. However, we show for the first time in macrophages that AMPK activation via A-769662 results in an up-regulation of ULK1 and Beclin-1 protein levels. This was associated with increased TFEB-regulated gene expression as well as increased TFEB nuclear localization. Furthermore, given that atherogenic lipoproteins activated AMPK, which in turn can influence TFEB-regulated pathways, it is possible that AMPK mediates part of the compensatory induction of lysosomal biogenesis by atherogenic lipids reported by others previously^34^.

We predicted that atherogenic lipoproteins would regulate transcriptional expression of lysosomal-associated genes through AMPK activation. However, our results were complicated by the fact that neither agLDL nor oxLDL had any effect on TFEB-associated transcript expression in WT macrophages, which had been shown previously^34^. Additionally, and in contrast to basal treatment without lipids, the expression levels in AMPKβ1-deficient cells treated with oxLDL were at times opposite. While there could be batch-to-batch and technical/experimental variation in preparing oxLDL (also in-house preparation vs. commercial suppliers), the type of macrophage presents the biggest discrepancy between ours and published results^34^. Thioglycolate-elicited peritoneal macrophages are a heterogeneous population of infiltrating monocyte-derived macrophages, which are often considered more activated in comparison to the homogenous population of bone marrow-derived cells^49^. The metabolic and inflammatory characteristics of these two primary murine macrophages have been investigated and could provide an explanation as to the discrepancy. Future work using elicited peritoneal and bone marrow-derived cells from WT and AMPKβ1-null mice would help bridge the divide between results.

Our data support that physiological, modified and atherogenic lipoproteins activate macrophage AMPK signaling via changes in calcium homeostasis. The activation of macrophage AMPK resulted in an up-regulation of lysosomal and autophagic pathways, responses that were blunted or not observed in AMPKβ1-deficient macrophages. Although endogenous AMPK signaling was important for various aspects of lysosomal homeostasis, in the presence of atherogenic lipoproteins, AMPK activation was insufficient to augment the same lysosomal programs. In the context of progressing of atherosclerosis, there could be a futile cycle, whereby plaque-resident macrophages are continually exposed to atherogenic stimuli that activate AMPK/TFEB related gene programs. However, the potential benefit stemming from acute AMPK activation, including the stimulation of autophagy and lysosomal pathways, are paradoxically not engaged. Rather, autophagy becomes progressively defective during the progression of atherosclerosis despite potential alterations in AMPK signaling^50^. Endogenous regulation of AMPK by pro-atherogenic lipoproteins does not rescue defective autophagy in cultured atherosclerotic macrophages, however whether pharmacological activators of AMPK are sufficient to overcome this hurdle remains to be tested.

## MATERIAL AND METHODS

### Animals

The generation and characterization of the AMPKβ1 knockout mice have been previously described^51^. Mice were generated from heterozygous pairings such that control mice were littermates. Mice were maintained on a 12-h light/dark cycle (lights on at 7:00 a.m.) and housed at 23 °C with bedding enrichment in a ventilated cage system. Male and female mice ages 8–16 weeks were used for the generation of primary macrophages as described below. CD36-deficient cells were isolated from the femurs of CD36 heterozygous mice, a kind gift from Dr. Kathryn Moore at NYU. All animal procedures were approved by the University of Ottawa Animal Care Committee.

### Isolation and culturing BMDM

Mice were anesthetized with ketamine (150 mg/kg) and xylazine (10 mg/kg), then euthanized by cervical dislocation. Each hind leg was carefully excised from the mouse then the legs were cleaned of soft tissue unlit bones were completely devoid of all tissue. both ends of the femur and tibia were carefully cut, bones were then placed a 0.5 mL tube containing a hole punctured at the bottom with 18-gauge needle at the bottom prior. The 0.5 mL tube containing bones was placed in a sterile 1.5 mL microfuge tube with 100 μL of culture media; Dulbecco’s modified Eagle’s medium (DMEM) supplemented with 10% fetal bovine serum and 100 U/mL Penicillin-Streptomycin (culture media). The tubes were centrifuged on a benchtop microcentrifuge at 4000 RPM for 5 min. The bones were discarded and 900 μL of culture media was added to each tube containing the bone marrow pellets, the pellets were gently re-suspended with gentle pipetting and then strained through a 40 μm cell strainer into culture media. Bone-marrow containing media was supplemented with 15% L929-conditioned media (and additional culture media) and plated within 150 mm tissue culture plates as needed. Cells were placed in a humidified incubator (37°C with 5% CO_2_) for 7-8 days to allow for macrophage differentiation. Following, media was removed from each 150 mm plate, cells were washed once with PBS, and then gently scrapped in culture media. Cells were counted and seeded as necessary to meet experimental

### Protein signaling experiments

BMDM were harvested and plated within a 6-well tissue culture plate at a density of 1.2 million cells per well (duplicate wells plated for each experimental condition) in culture media. All experiments began with removal of culture media, cells were washed once with ice-cold PBS. Following the PBS was removed and 1 mL of fresh culture media was added to each well with experimental treatments. Experiments were conducted such that the longest time-points (if applicable) were treated first to ensure all conditions were finished simultaneously. At experimental endpoint, all media was removed and cells were washed with PBS. PBS was removed and cells were scraped in 80 μL lysis buffer (50 mM Tris.HCl pH 7.5, 150 mM NaCl, 1 mM EDTA, 0.5% Triton X-100, 0.5% NP-40, 100 μM Na_3_VO_4_, supplemented with protease inhibitor cocktail (PIC) tablet. Duplicate lysates were combined and placed in a new 1.5 mL microfuge tube and snap frozen with liquid nitrogen. Lysates were thawed on ice, following samples were centrifugation at 14000 RPM at 4°C for 5 m. The pellet was discarded and the lysate was collected and placed in a new 1.5 mL and stored at −80°C.

### Protein quantification, normalization, and preparation

Total protein quantification was determined using the Pierce™ BCA Protein Assay Kit as per manufacturers instructions under the microplate procedure and reading samples with Biotek’s Gen5 software. Following quantification, protein samples were normalized to the 1.34 μg/μL in lysis buffer, then an aliquot was further diluted in 4X loading dye; 200 mM Tris-HCl pH 6.8, 400 mM Dithiothreitol (DTT), 8% Sodium dodecyl sulfate, 0.4% bromophenol blue, 40% glycerol. Protein samples in loading dye were then boiled at 95°C for 5 m. Samples were stored at −20°C until ready for analysis.

### LDL aggregation

1 mL of 5 mg/mL LDL was vortexed at max speed for precisely 60 seconds.

### Immunoblotting

Equalized protein samples containing loading dye were loaded (15 μg protein/well) and electrophoresed onto either a 4-20% or 8% SDS-PAGE gel (110 volts for ~1.75 h). For analysis of relative phosphorylation of target proteins, duplicate gels were electrophoresed in tandem to probe for phosphorylated and total levels for target proteins. Following electrophoresis, each gel was carefully removed and transferred onto PVDF membranes using Bio-Rad’s Trans-Blot Turbo system (25 V, 2.5 amps, for 18 min). Membranes were then blocked in TBST supplemented with 5% w/v BSA with gentle rocking at room temperature for 1 h. Membranes were then cut at appropriate molecular weights to isolate multiple proteins of interest per membrane. Each membrane fraction was incubated in primary antibody solution (TBST supplemented with 5% BSA and 1:1000 dilution of primary antibody of interest overnight at 4°C. The following day the primary antibody solution was collected and membranes were washed four sequential times in TBST for 5 min with gentle rocking at room temperature. After washing, membranes were incubated with anti-rabbit HRP conjugated secondary antibody (TBST supplemented with 5% BSA and 1:10000 dilution of anti-rabbit secondary) for 1h at room temperature with gentle rocking. Following, secondary-antibody solution was discarded and membranes were washed four times as previously done. Image acquisition was completed by gently drying membranes then incubating briefly in Clarity ECL substrate mix. Luminescence was measured using the General Electric’s ImageQuant LAS4000 system, while obtained image files were processed and quantified using Image J analysis software.

### Relative transcript expression experiment

BMDM were harvested and plated within a 12-well tissue culture plate at a density of 0.6 million cells per well (triplicate wells plated for each experimental condition) in culture media. All experiments began with removal of culture media, cells were washed once with ice-cold PBS. Following the PBS was removed and 0.5 mL of fresh culture media was added to each well with experimental treatments. Experiments were conducted such that the longest time-points (if applicable) were treated first to ensure all conditions were finished simultaneously. At experimental endpoint, all media was removed and 0.5 mL of TriPure reagent was added directly to the cells. TriPure containing plates were flash frozen by incubating the plate in a Styrofoam container with a shallow level of liquid nitrogen. Plates were thawed on ice, then samples in TriPure were collected following visual confirmation that adherent cells are not present (ensuring proper lysis) and placed within a sterile 1.5 mL microfuge tube (replicates were collected separately). Samples were then either stored at −80°C or were subjected to an RNA isolation (explained below).

### RNA Isolation

RNA isolations were performed using Roche’s TriPure reagent kit as per manufacturers instructions. Following isolation, the RNA pellet was then resuspended in 30 μL of nuclease free water (Wisent) and incubated at 58°C for 15 m. Samples were stored at −80°C until quantification and synthesis of cDNA were ready to be performed.

### RNA quantification

RNA concentration and purity were assessed using a take-3 plate reader. In brief, 2 μL of sample was placed on corresponding location within the take-3 plate in duplicate using nuclease free water as a blank. Samples were then measured using Gen5 software, samples with a 260/280 absorbance ratio less than 1.8 or greater than 2.0 were removed from assessment. All samples were then normalized to 20 ng/mL in nuclease free water, now ready for cDNA synthesis.

### cDNA synthesis

Genomic DNA was removed from all RNA samples using AccuRT Genomic DNA removal kit as per manufacturers instructions. Following cDNA synthesis from RNA was performed using Applied Biological Material’s 5X All-In-One RT Master mix product as per manufacturers instructions. Following cDNA synthesis, samples were further diluted 1:20 with nuclease-free water prior to transcript expression analysis.

### Quantitative PCR

Quantitative PCR reactions were performed within 0.1 mL 4-Strip tubes using 4.75 μL of diluted cDNA template, 0.25 μL of the TaqMan primer/probe set of interest, and 5 μL of PCR master mix and run on a Rotorgene Q in a two-step amplification reaction.

### Immunofluorescent labeling

Immunofluorescent experiments were conducted on 8-well chamber slides, each chamber was seeded with 5 x 10^4^ BMDM. Upon experimental completion, all media was removed and each chamber slide was washed once with PBS. PBS was removed, and cells were fixed for 20 min with 300 μL of 2% PFA at room temperature. The fixative was removed and chambers were washed twice with PBS. PBS was removed and fixed cells were blocked/permeabilized for 1 h at room temperature with PBS supplemented with 300 μL PBS supplemented with 5% fatty-acid free BSA, 0.2% Triton X-100 and 0.1% Tween-20. Blocking/permeabilizing buffer was removed, chambers were washed once with PBS, then 300 μL of anti-TFEB antibody solution (PBS supplemented with 1:100 anti-TFEB primary antibody, 2% fatty acid-free BSA, 0.2% Triton X-100, 0.1% Tween-20). All chamber slides were incubated in primary antibody solution over night at 4°C. The following day, the antibody solution was removed, chambers were washed twice with PBS. PBS was removed and chamber slides were incubated in 300 μL secondary antibody solution (PBS supplemented with 1:100 anti-mouse Alexa 488-conjugated secondary antibody, 2% fatty acid-free BSA, 0.2% Triton X-100, 0.1% Tween-20) for 1 h at room temperature shield from light. Following, secondary antibodies solution was removed and chambers were washed once with PBS. PBS was removed and 300 μL of 300 nM DAPI made in PBS was added to each chamber and let incubate temperature for 5 m shielded from light. Following, DAPI solution was removed and all chambers were washed three times with PBS. Chambers were stored in 300 μL PBS at 4°C shielded from light until image acquisition was ready to be performed.

### Immunofluorescent imaging and nuclear colocalization

Following immunolabeling, multiple Z-stack images spanning the entirety of the cell at 50 μm increments were taken at 20X using a Zeiss LSM800 AxioObserver Z1 Confocal microscope. Image files were processed using Image J Software and representative Z-projection for maximum intensities were used for representative images. Colocalization between DAPI and TFEB channels throughout all z-stacks (23 slices) were performed using the EZColocalization plugin on Image J software as reported^52^.

### AMP and ATP determination

BMDM were harvested and plated within a 100 mm tissue culture plate (Thermo Fisher Scientific) at a density of 5 million cells per plate in culture media (duplicate plates were seeded for each experimental condition). At the experimental endpoint, media was removed from each plate and cells were washed once with ice-cold PBS. Following, 250 μL ice-cold perchloric acid (10% v/v HClO_4_, 25 mM EDTA) was added and cells were scraped/lysed with a rubber policeman. Cellular extracts were transferred to a sterile 1.5 mL microfuge tube and vortexed briefly. Samples were left to incubate on ice for 30 min then precipitated proteins were spun down at 8000 *g* for 2 min at 4°C. A 200 μL aliquot was taken and placed in a new microfuge tube, samples pH was adjusted to 6.5-7 using ~130 μL KOH/MOPS (2 N KOH/0.3 M 3-N-morpholino propane sulfonic acid) with careful mixing and confirmation using pH strips. Lysate was clarified by centrifugation at 8000 x *g* for 2 min at 4°C. A 200 μL aliquot of neutralized supernatant fraction was pipetted into a new 1.5 mL microfuge tube and was stored at −80°C until ready for high performance liquid chromatography measurement (HPLC). HPLC measurements were performed as described^53^.

### Reagents

4’,6-Diamidino-2-Phenylindole, 2-(4-amidinophenyl)-1H -indole-6-carboxamidine (DAPI; Thermo Fisher Scientific – D1306), 5X All-In-One RT MasterMix with AccuRT Genomic DNA Removal Kit (Applied Biological Material - G492), A-769662 (AdooQ Bioscience – 844499-71-4), Acetylated LDL (Alfa Aesar - J65029), Albumin (Albumin, Bovine, Serum, Heat Shock Isolation, Fraction V. Min. 98%; Bioshop - ALB001.250), BCA™ Protein Assay (Thermo Fisher Scientific - PI-23225), Butylated hydroxyanisole (Sigma-Aldrich - B1235-5G), Chloroform, ACS, Reagent Grade, 1 L (Bioshop Canada - CCL402.1), Chloroquine (Sigma-Aldrich - C6628-25G), Clarity Western ECL Substrate (Bio-Rad - 170-5060), Complete Protease Inhibitor Cocktail Tablets Mini, EDTA-Free - 30 units (Sigma-Aldrich - 4693159001), CytoPainter Lysosomal Staining Kit - Blue Fluorescence (Abcam - ab112135), Dextran, Tetramethylrhodamine, 10,000 MW, Lysine Fixable (Thermo Fisher Scientific - D1817), DFQ™ Ovalbumin (Thermo Fisher Scientific - D12053), Dithiothreitol (DTT), Electrophoresis Grade (Bioshop Canada - DTT001.10), DMEM (Wisent – 319-005-CL), DMSO (Thermo Fisher Scientific – 23730571), EDTA (Wisent - 625-060-CG), Ethanol (Commercial Alcohols - P016EAAN), Fatty Acid-Free Powder (Thermo Fisher Scientific – BP9704100), Fetal bovine serum (Wisent – 095-150), Hydrochloric Acid (HCl) (Thermo Fisher Scientific – 8732113), Isopropanol (Thermo Fisher Scientific - A416-4), LDL (Alfa Aesar - J65039), Lipofectamine™ RNAiMAX Transfection Reagent (Thermo Fisher Scientific - 13778075), MOPS (Sigma-Aldrich - M3183-25G), NP-40 (Bioshop Canada - NON999.500), Oligomycin A (Cedarlane - 4110 n/5), Opti-MEM Reduced Serum Medium (Invitrogen - 31985-070), Oxidized LDL (Alfa Aesar - J65591), Paraformaldehyde (Bioshop Canada – PAR070.500), PBS (Wisent – 311-010-CL), Penicillin/Streptomycin (Thermo Fisher Scientific −SV30010), Perchloric Acid ACS reagent 70% (Sigma-Aldrich - 311421-250mL), Potassium Hydroxide, Reagent Grade (Bioshop Canada - PHY202.500), ProLong™ Gold Antifade Mountant (Thermo Fisher Scientific - P36930), Ryanodine (Tocris - 13-291), Sodium Chloride (Bioshop Canada - SOD001.1), Sodium Orthovanadate (Bioshop Canada - SOV664.10), STO-609-acetic acid (Abcam - ab141591), TriPure (Roche - 11667165001), Tris (Bio Basic - TB0196), Triton X-100 (Bioshop Canada – TRX777.500), Tunicamycin (Sigma-Aldrich - T7765), Tween-20 (Thermo Fisher Scientific – BP377-500), U18666A (Sigma-Alrich - U3633), Water Rnase and Dnase free (Wisent - 809-115-CL).

### Antibodies

Acetyl-CoA Carboxylase (Cell Signaling Technology - 3676), AMPKα (Cell Signaling Technology - 5831), Anti-Mouse CD107a (LAMP-1) eFluor 450 (Cedarlane - 48-1071-82), Anti-rabbit IgG HRP-linked Antibody (Cell Signaling – 7074S), ATF-6 (Cell Signaling Technology - 65880), Beclin-1 (Cell Signaling Technology - 3495), Beta-Actin (Cell Signaling Technology - 5125), CHOP (Cell Signaling Technology - 2895), LC3A/B (D3U4C) (Cell Signaling Technology - 12741), mTOR (Cell Signaling Technology - 2983), Phospho-Acetyl-CoA Carboxylase Ser79 (Cell Signaling Technology - 11818), Phospho-AMPKα Thr172 (Cell Signaling Technology - 2535), Phospho-mTOR Ser2481 (Cell Signaling Technology - 2974), Phospho-Raptor Ser792 (Cell Signaling Technology - 2083), Phospho-Tuberin/TSC2 Ser1387 (Cell Signaling Technology - 23402), Phospho-ULK1 Ser555 (Cell Signaling Technology - 5869), Phospho-ULK1 Ser757 (Cell Signaling Technology - 6888), Raptor (Cell Signaling Technology - 2280), Rat IgG2a Isotype Control eFluor 450 (Cedarlane - 48-4321-80), SQSTM1/p62 (Cell Signaling Technology - 5114), TFEB, Monoclonal Antibody (Mybiosource, inc - MBS120432), Tuberin/TSC2 (D93F12) (Cell Signaling Technology - 4308), ULK1 (Cell Signaling Technology - 8054).

### Statistical analyses

All statistical analyses were completed using GraphPad Prism 7.03 (GraphPad Software Inc.). Comparison between two groups were made using a paired, two-tailed Student’s t-test. Comparisons between more than two treatment groups were made using a one-way ANOVA analysis with a Tukey test for multiple comparisons. For experiments comprising more than two groups (genotype and treatment), a two-way ANOVA was used with a Tukey test for multiple comparisons. Significant differences are described in the figure legends. All data are expressed as mean±SEM, unless specified in the figure legend.

## Acknowledgements

We would like to thank Dr. Chloë van Oostende-Triplet and Skye McBride in the Cell Biology and Image Acquisition Core, as well as Dr. Vera Tang in the Flow Cytometry Core for technical assistance. We would also like to thank Dr. Kathryn Moore (NYU) for CD36-deficient femurs.

## Supplementary Figure Legend

**Supplementary Figure I**. Atherogenic lipoproteins agLDL and oxLDL enhance AMPK-specific signaling to ACC. (A) Representative immunoblot depicting increased AMPK signaling in response to incubation with agLDL (50 μg/mL) for 8 and 24 h. A-769662 (8h; 100 μM) was used as a positive control for AMPK activation. (B) Representative immunoblot depicting increased AMPK signaling in response to incubation with oxLDL (50 μg/mL) for 8 and 24 h. A-769662 (8h; 100 μM) was used as a positive control for AMPK activation. (C) Quantification of relative signaling to ACC at Ser79 in response to 8 and 24 h incubation with atherogenic lipoproteins agLDL and oxLDL. Data represent the mean ± SEM.

**Supplementary Figure II**. Atherogenic lipoproteins uncouple macrophage AMPK’s ability to regulate lysosomal-associated gene transcription. (A-C) Relative transcript expression for (A) *Tfeb*, (B) *Beclin-1* and *Lipa* (lysosomal acid lipase) in the presence of atherogenic lipoproteins agLDL and oxLDL (50 μg/mL) over time. Transcript expression was made relative to untreated WT macrophages and normalized to the average of *β actin* and *Tbp*. Data represent the mean ± SEM.

