## Supplementary figures and images for "Foam cell induction activates AMPK but uncouples its regulation of autophagy and lysosomal homeostasis"

### Supplementary Figure S1

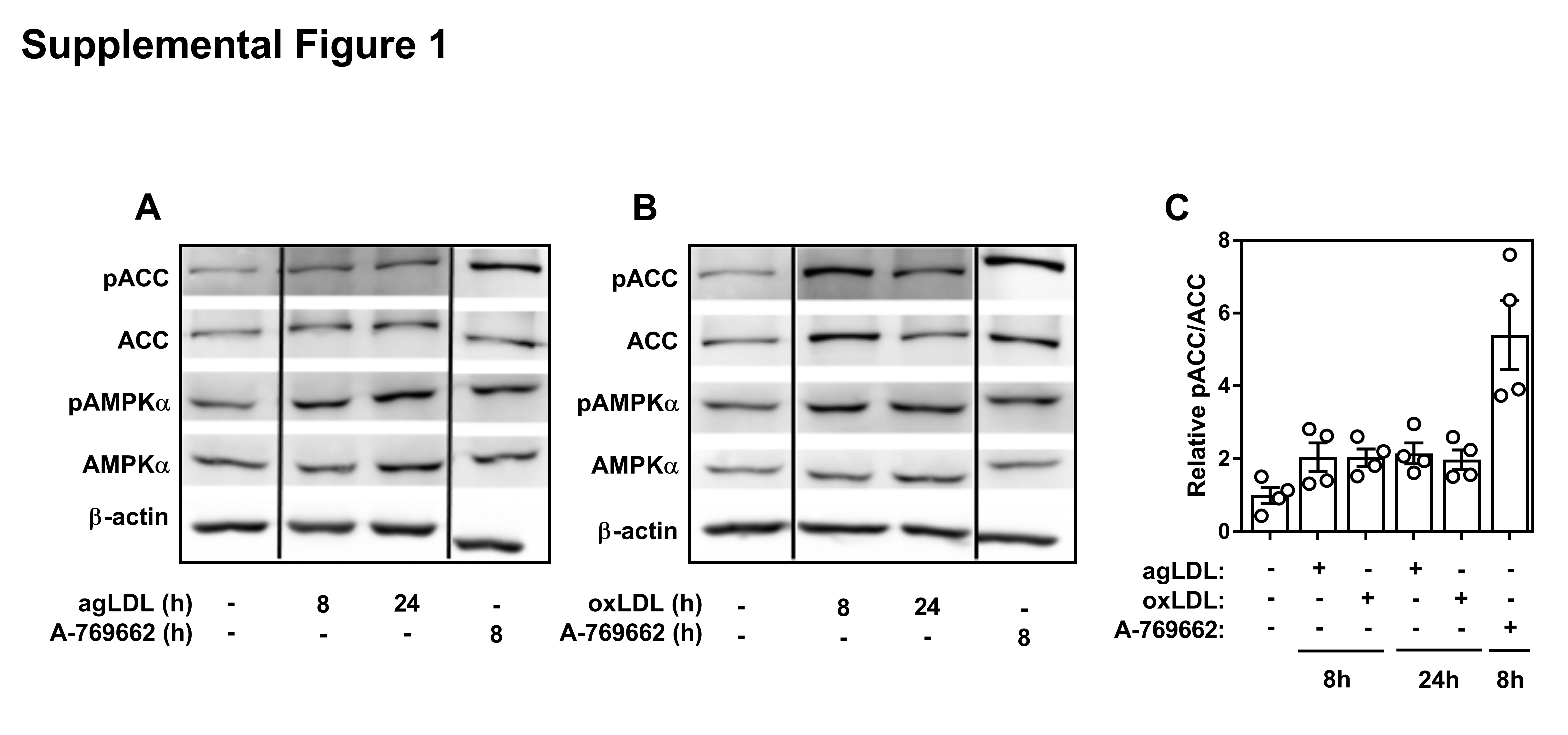

### Supplementary Figure S2

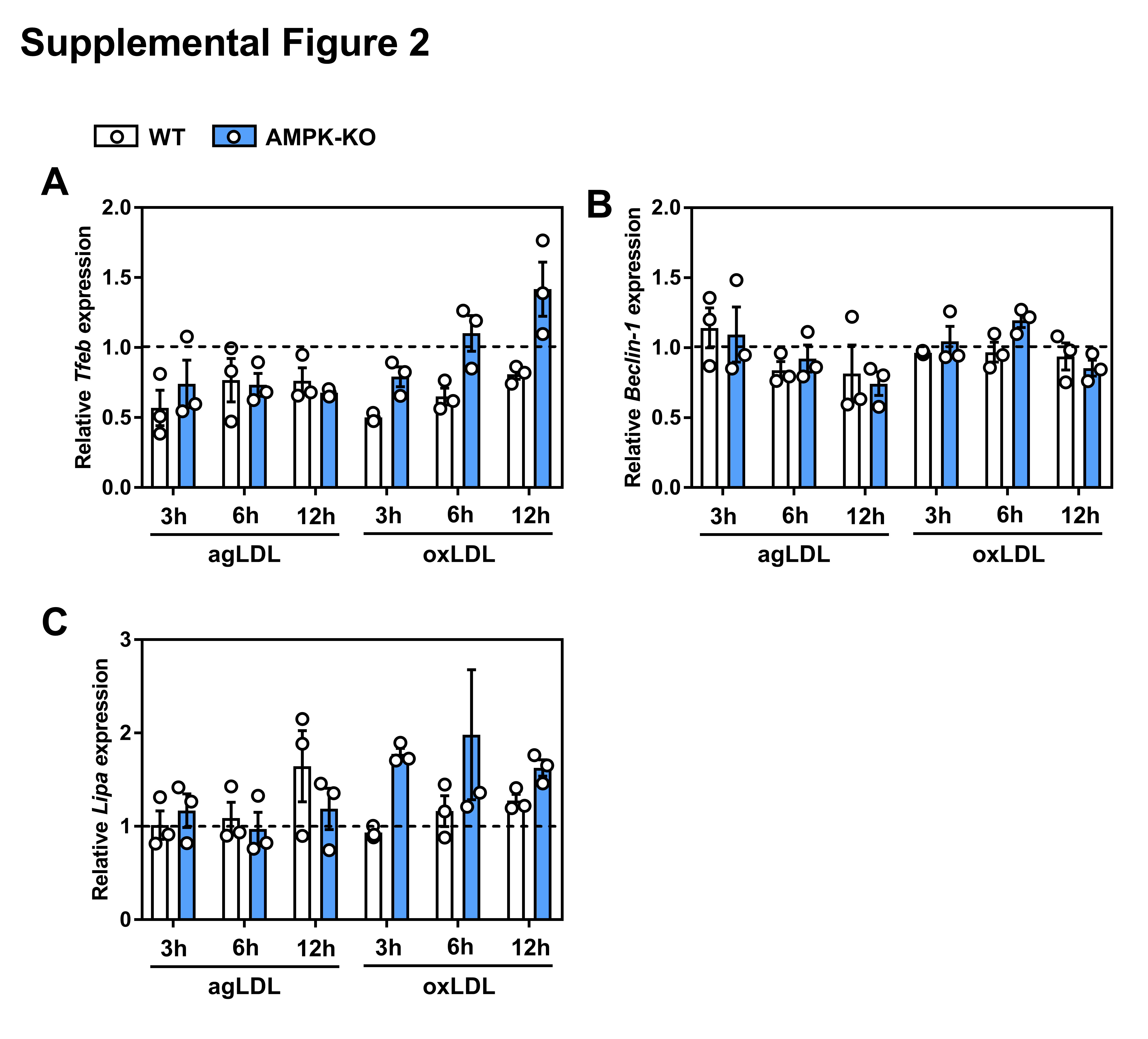
